# Versatile Multi-Detector Scheme for Adaptive Optics Scanning Laser Ophthalmoscopy

**DOI:** 10.1101/408526

**Authors:** Sanam Mozaffari, Volker Jaedicke, Francesco Larocca, Pavan Tiruveedhula, Austin Roorda

**Author notes:** Contributed equally to this work (Sanam Mozaffari, Volker Jaedicke).

## Abstract

Adaptive Optics Scanning Laser Ophthalmoscopy (AOSLO) is a powerful tool for imaging the retina at high spatial and temporal resolution. In this paper, we present a multi-detector scheme for AOSLO which has two main configurations: pixel reassignment and offset aperture imaging. In this detection scheme, the single element detector of the standard AOSLO is replaced by a fiber bundle which couples the detected light into multiple detectors. The pixel reassignment configuration allows for more light throughput while maintaining optimal confocal resolution. The increase in signal-to-noise ratio (SNR) from this configuration can improve the accuracy of motion registration techniques. The offset aperture imaging configuration enhances the detection of multiply scattered light, which improves the contrast of retinal vasculature and inner retinal layers similar to methods such as nonconfocal split-detector imaging and multi-offset aperture imaging.

## 1. Introduction

Adaptive Optics Scanning Laser Ophthalmoscopy (AOSLO) is a powerful retinal imaging tool with high spatial and temporal resolution [1]. It allows for the imaging of retinal structures at a cellular scale, such as cone photoreceptors, at localized positions within retinal tissue. An AOSLO is a confocal scanning laser microscope that incorporates adaptive optics and utilizes the eye’s optics as the microscope objective. Just as in confocal microscopy, a focused spot is scanned across the sample (the human retina) and a single element integrating detector captures the backscattered light at every scan position. A pinhole is placed in front of the detector at a plane conjugate to the focused spot on the sample to reject out-of-focus light. The diameter of this pinhole is one of the key elements that governs the resolution of the microscope [2]. The lateral resolution is improved by a factor of 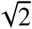 over a widefield flood illumination instrument when a infinitesimally small pinhole is introduced. Increasing the pinhole size allows for more light throughput at the cost of decreasing the resolution. Due to this trade off between signal and resolution, the common pinhole size for AOSLO is generally designed to fall between 0.5*ADD* and 1*ADD* [3, 4], where ADD refers to the Airy Disc Diameter, determined by *ADD* = 2.44 · *λ · f/D*, where *f* is the focal length of the collector lens, *λ* is the wavelength of light and *D* is the beam diameter.

In this paper, the single element integrating detector is replaced by a multi-detector consisting of a bundle of 7 multi-mode fibers in the form of a hexagonal array. Each fiber is directly coupled to a photomultiplier tube enabling 7 simultaneous image acquisition channels. Multi-detection schemes offer the ability to extend imaging of scanning systems beyond the traditional confocal mode.

One way to employ multi-detector imaging is pixel reassignment, in which images from individual detectors are registered and added [5]. The resolution of the final image is governed by the size of one detector element as in a standard confocal microscope [5, 6], but by combining the signal from multiple detectors, the system throughput is increased. Pixel reassignment methods have been successfully applied to improve resolution and SNR in confocal fluorescence microscopy using detector arrays [7–10] or cameras [11–14]. In order to improve the acquisition speed, camera-based methods have also been combined with multi-spot excitation [15]. Alternatives to this digital post-processing approach that employ a camera with analog, all-optical processing have also been demonstrated [12–14, 16, 17] and recently implemented for a scanning laser ophthalmoscope without adaptive optics [18]. However, the reduced system complexity for this all-optical processing comes with the price of less flexibility in terms of data post-processing.

A multi-detector scheme can also be reconfigured to facilitate the collection of multiply-scattered, and refracted light, which has proven to have great utility for retinal imaging. Detection of non-confocal, multiply-scattered light has been used to reveal subretinal structures [19] and retinal pigmented epithelium cells in the human retina [20]. Asymmetric, or offset aperture detection schemes (i.e collecting and/or comparing multiply-scattered light from different directions relative to the confocal aperture) have revealed transparent and/or refracting structures in the retina including photoreceptor inner segments [21], blood vessels [22], blood cells [23], horizontal cells [24] and ganglion cells [25].

The parallel nature of multi-detector schemes offer increased efficiency, flexibility and fidelity over single acquisition [22], or multiple serial acquisition [25] techniques for offset aperture detector imaging. In the earliest implementation, simultaneous collection of two spatially offset channels was achieved by using a reflective mask to separate nonconfocal and confocal light; the light on either side outside a central area being transmitted through a mask and collected by two nonconfocal detectors, while the light reflected off the center of the mask was directed to a confocal detector [21]. More recently, this approach with a static mask has been further modified using a programmable, pixelated reflector to direct the light outside the confocal aperture into two detectors [26] which allowed for the use of arbitrary aperture shapes and orientations for vessel imaging.

The multi-detector scheme described in this paper can be configured to offer both pixel reassignment and multi-offset aperture imaging modalities. In the results section, we describe how the system is setup, then we show imaging results from healthy human volunteers. Finally, we discuss further applications and improvements in the discussion section.

## 2. Methods

### 2.1. AOSLO System with Multi-detector

The specific multi-wavelength AOSLO system is described in more detail in previous publications [27] and so only the details that are most pertinent to this paper are described here. Only the 680 nm imaging channel was modified in the multi-wavelength AOSLO for the multi-detector scheme. A system schematic is shown on the left of figure 1.

**Fig. 1.**
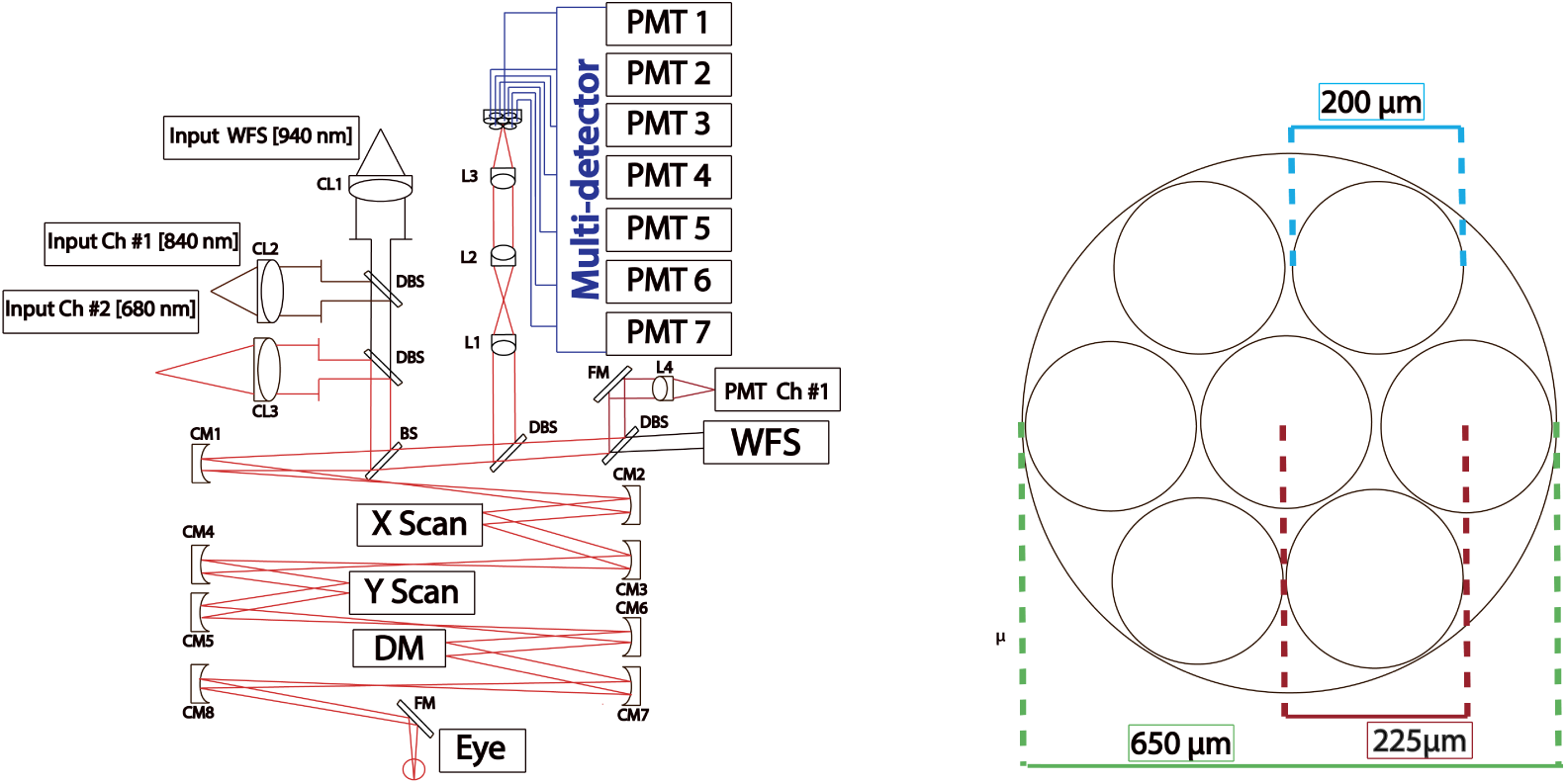
Schematic of the AOSLO system with a 840 nm PMT channel and a 680 nm multi-detector channel. The key component in the multi-detector setup is the 4f telescope (L2-L3) which relays the image of the focused spot on the retina to the fiber core. The geometry of the multi-detector’s fiber bundle is shown on the right. (*BS*: *Beamsplitter, CL*: *Collimating Lens, CM: CurvedMirror, L*: *Lens, DBS*: *DichroicBeamsplitter, DM*: *De f ormableMirror, PMT*: *PhotomultiplierTube, WFS*: *Wave frontSensor*)

**Fig. 2.**
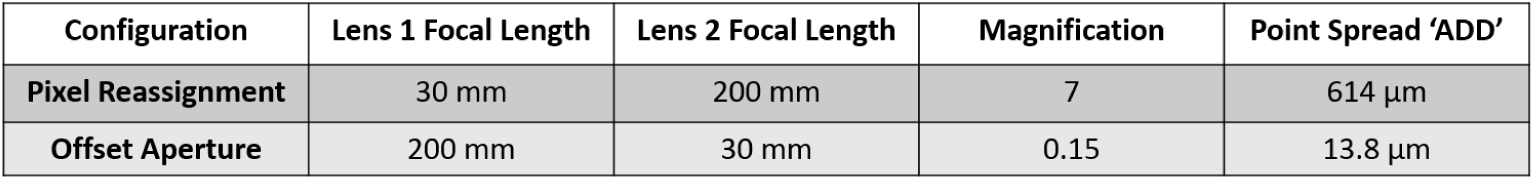
Summary of the optical configuration for the desired point spread for each multi-detector configuration.

The system’s optical design was modeled after the AOSLO design described by Dubra et al. [4]. The front end (double-pass part of the system comprised of the optics between the first beam splitter (BS) and the eye) consists of afocal telescopes, formed by pairs of off-axis spherical mirrors in a non-planar arrangement. The eye’s pupil plane is imaged onto a deformable mirror (DM97-08, ALPAO, Montbonnot-Saint-Martin, France), a galvo scanner (6210h, Cambridge Technology, Bedford, USA) and a resonant scanner (SC-30, EOPC, Ridgewood, USA). The deformable mirror is the pupil stop with a diameter of 7.2*mm*. Light from a supercontinuum laser (SuperK Extreme, NKT Photonics, Birkerod, Denmark) is bandpass filtered such that it has a central wavelength of 680*nm* (22*nm* bandwidth) and 840*nm* (22*nm* bandwidth) for imaging and 940*nm* (10*nm* bandwidth) for wavefront sensing. All wavelength bands are combined using a dichroic beamsplitter (DBS) and reflected off a 0.5 degree wedge 10/90 (R:T) beam splitter (BS). After passing through the front end, light backscattered from the retina is descanned and transmitted through the beam splitter into the collection optics. In the case of the multi-detector, the retinal conjugate spot formed by Lens 1 is relayed by a 4f telescope (Lens 2 and Lens 3) onto a multi-mode fiber bundle (BF72HS01, Thorlabs, Newton, USA) in which the fibers are arranged in a closely packed hexagonal array and then split into seven individual fibers. The geometry of the common end of the fiber bundle is given on the right side of figure 1. The diameter of each individual fiber is 200*µm*, while the fiber pitch is 225*µm*. The overall diameter of the fiber bundle tip is 650*µm*, which corresponds to a fill factor of 66%.

### 2.2. Multi-Detector Telescope

In the original configuration, Lens 1 was the collector lens that focused the 680 nm light to a single confocal pinhole. With a beam size of 3.6 mm and a focal length of 200 mm, the ADD of the focused spot was 92*µm*.

A 4f telescope was added to the system to relay an image of that spot onto the fiber bundle. The focal lengths of the lenses in the telescope determine the imaging configuration of the multi-detector since they control the size of the focused spot relative to the fiber bundle core. For pixel reassignment mode, Lens 2 and Lens 3 were set to *f* = 30*mm* and *f* = 200*mm* (AC254B, Thorlabs, Newton, USA), respectively, to provide a magnification factor of 7, a collection spot size of 614*µm*, and a ratio of the fiber core to PSF diameter of 0.33*ADD*. The entire collection diameter of the fiber bundle was 1.07*ADD* as shown in Figure 3.

**Fig. 3.**
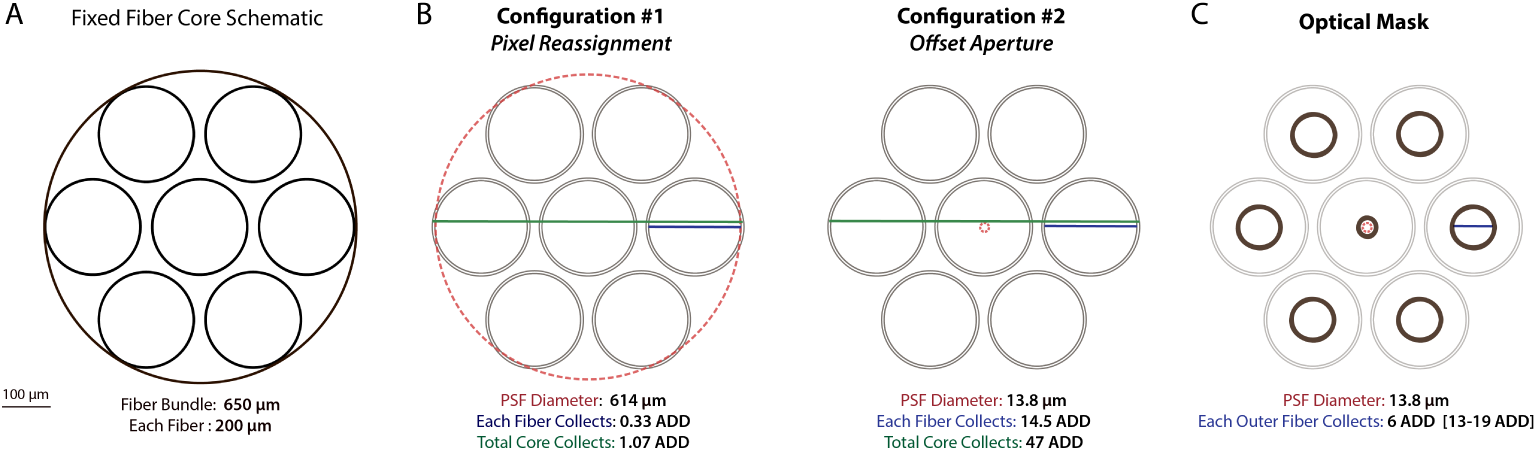
A. Geometry of fiber bundle. B. For pixel reassignment, the collection PSF is magnified by 7 to be the same size of the fiber core, 614*µm*. For offset aperture imaging, the collection PSF is magnified by 0.15 to be smaller than the fiber core, 13.8*µm*. C. Optical Mask for offset aperture imaging to limit the light collection from 13 to 19 ADD with 6 ADD apertures for each fiber

For offset aperture imaging, the two lenses in the telescope were merely flipped such that the first lens of the telescope was *f* = 200*mm*, while the second was *f* = 30*mm* for a magnification of 0.15 resulting in a PSF diameter of 13.8*µm* and a fiber core-to-PSF diameter ratio of 14.5*ADD*. The entire collection area of the fiber bundle was 47*ADD* in this mode. Additionally, an optical mask was used to collect over a 6*ADD* diameter aperture for the outer fibers and provide a 2*ADD* confocal pinhole for the central fiber. The optical mask was inserted one focal length away from Lens 1 and mounted on a three axis kinematic stage with additional rotation adjustment.

### 2.3. Acquisition and Data Processing

The common end of the fiber bundle was mounted on a linear kinematic stage (PI 403.8DG, Physik Instrumente, Karlsruhe, Germany), which enabled movement of the detector along the optical axis. Each individual fiber was connected to a PMT (Photo Multiplier Tube). Since 7 identical PMTs were not available, different PMT models (7422-20, 7422-40 and 7422-50, Hamamatsu, Hamamatsu, Japan) which had similar performance at 680*nm* were fiber coupled to the multi-detector. Each of the PMT modules was equipped with an individual gain control to adjust for small differences in performance and light levels at different detection positions. Each channel’s signal was further amplified using identical amplifiers (C6438-01, Hamamatsu, Hamamatsu, Japan) for each PMT. Custom electronics were used to correct the black level of the signal, apply a temporal apodization window, and low-pass filter the signal with a cutoff frequency of 10*MHz*. Two identical 4-channel framegrabbers (Helios, Matrox Imaging, Dorval, Canada) were used for digitization of the signal and were triggered by a sync signal from the fast scanner. Custom acquisition software allowed for the simultaneous display of up to 8 channels. Frames were acquired at 30 frames per second with pixel dimensions of 512×512 pixels at a 0.8 ° field of view. The scanners were driven with custom electronics and custom software. Adaptive optics correction was based on a custom Shack-Hartman wavefront sensor and controlled using custom software.

All data processing was done in MATLAB (MATLAB, Mathworks Inc, Natick, USA) and ImageJ [28]. Raw videos from all seven channels were first corrected for non-linear sampling due to the sinusoidal waveform of the resonant scanner using calibration data obtained from a grid target. A strip-based stabilization software was used to estimate and correct for eye motion [29]. Frames that contained distortion from motion or blinks were removed and the average image intensity across all frames of the video was taken to form an image. Eye motion data obtained from the central detector was used to register all other acquired videos.

In pixel reassignment mode, images were registered with respect to the image acquired by the central detector. Individual images of the detector elements were displaced *d/* 2, where *d* is the geometrical distance with respect to the central detector element [30]. To account for small misalignment errors and the unknown rotation of the fiber bundle, shifts between different imaging channels were computed using a sub-pixel Fourier transform-based algorithm [31] instead of using theoretically determined values. The shifts for each imaging session were calculated from the best quality image set and subsequently applied to all images of the same data set.

For offset aperture imaging, the raw data was contrast enhanced to adjust for the different PMT response characteristics, histogram-matched to account for different PMT amplifier gains, and filtered with wavelet-based denoising to decrease image degradation due to noise. The differences of individual fiber images were taken according to the opposing positions and normalized with respect to their sum.

Due to the weak reflective signal within the inner retinal layers, neither the 840*nm* nor the 680*nm* imaging channel had enough SNR to register eye motion. In order to take a complete dataset through depth, the axial focal position of the 680*nm* beam and the conjugate multi-detector channel was offset to be anterior to the the 840*nm* imaging channel by about 200*µm* in the retina. In this way, the 840*nm* beam was focused on the photoreceptors which offered rich structure for optimal eye motion registration while the offset imaging channel was located just under the nerve fiber layer.

## 3. Results

Subjects: The University of California Berkeley Institutional Review Board approved this research, and subjects signed an informed consent before participation. All experimental procedures adhered to the tenets of the Declaration of Helsinki. Mydriasis and cycloplegia were achieved with 1% tropicamide and 2.5% phenylephrine ophthalmic solutions before each experimental session. Subjects bit into a dental impression mount affixed to an XYZ stage to hold the eye and head still. Both subjects were healthy young adult volunteers, 20112L and 20076R.

### 3.1. Pixel Reassignment

The multi-detector telescope was set up to magnify the focused spot by a factor of 7 to form a collection spot size of 614*µm* in front of the detector with a fiber core-to-PSF diameter ratio of 0.33*ADD*. The optical power output of the imaging system prior to the eye was measured to be 108.6*µW* for 940*nm* and 73.6*µW* for 680*nm*. First, 10-second videos (300 frames) from each individual detector were recorded and corrected for intra-frame eye motion and an average image was calculated.

In figure 4 images of the foveal center are shown. Each of the individual fiber images were collected with a 0.33 ADD pinhole. The Averaged Image from figure 4 was obtained by summing the images obtained from all detectors without applying the pixel reassignment process. In this case, the resolution and contrast was effectively equivalent in resolution and throughput to that obtained through a 1 ADD pinhole, aside from light losses due to the smaller fill factor of the fiber bundle (see discussion).

**Fig. 4.**
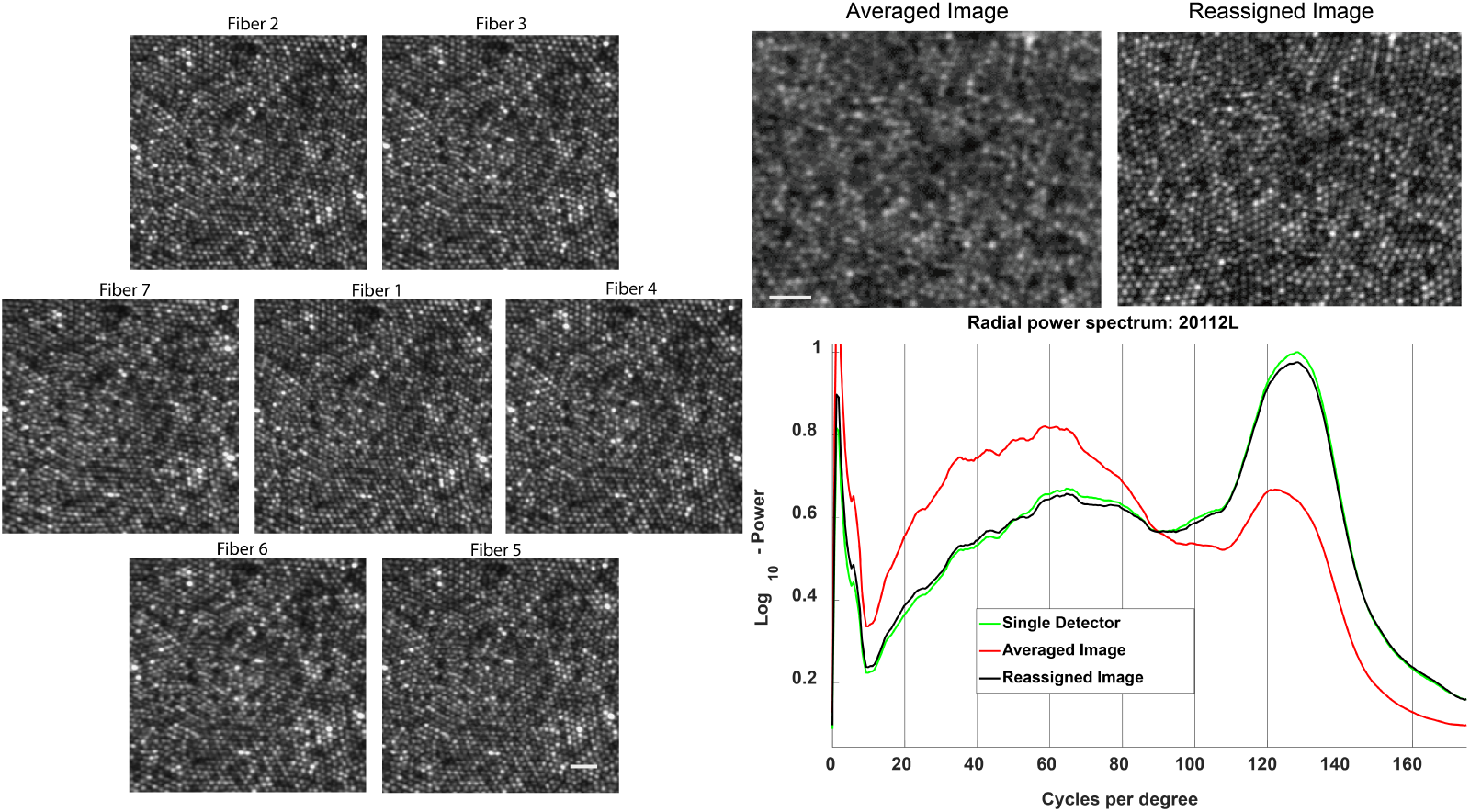
The hexagonal array of images (from Subject 20112) displays the images acquired with each fiber which collects approximately a 0.3 ADD of the collection PSF. The Averaged Image is a simple average of all 7 images from the hexagonal array of images shown on the left. In the Reassigned Image, the 6 outer images are registered to the central image prior to summation. The radial power spectrum quantifies the higher spatial frequency components in the image. The peak at 125 cycles per degree corresponds to the signal from the periodic cone mosaic. The power at the peak is similar to that of the image obtained via a single detector and the Reassigned Image, but is reduced in the power spectrum obtained from the Averaged Image. Scale bar: 10*µm*

The Reassigned Image shown in figure 4 is the sum of all 7 fibers after subpixel registration. The individual multi-detector images are almost indistinguishable in resolution from the Reassigned Image. The radial power spectrum in figure 4 shows that the high spatial frequencies in the single detector is preserved within the Reassigned Image unlike that of the Averaged Image. In other words, reassigned images collected through the multi-detector system allow for an increase in light collection without a compromise in image quality.

The benefits of increased light collection in pixel reassignment are illustrated in figure 5, where the superior performance of pixel reassignment is clearly visible. In panel A, an average of 5 frames from the multi-detector scheme was enough to virtually remove noise and motion artifacts, while the 5 frame average using the single detector scheme still visibly suffered from both. In other words, the use of a multi-detector with the pixel reassignment scheme allowed the acquisition of high SNR images in a shorter period of time.

**Fig. 5.**
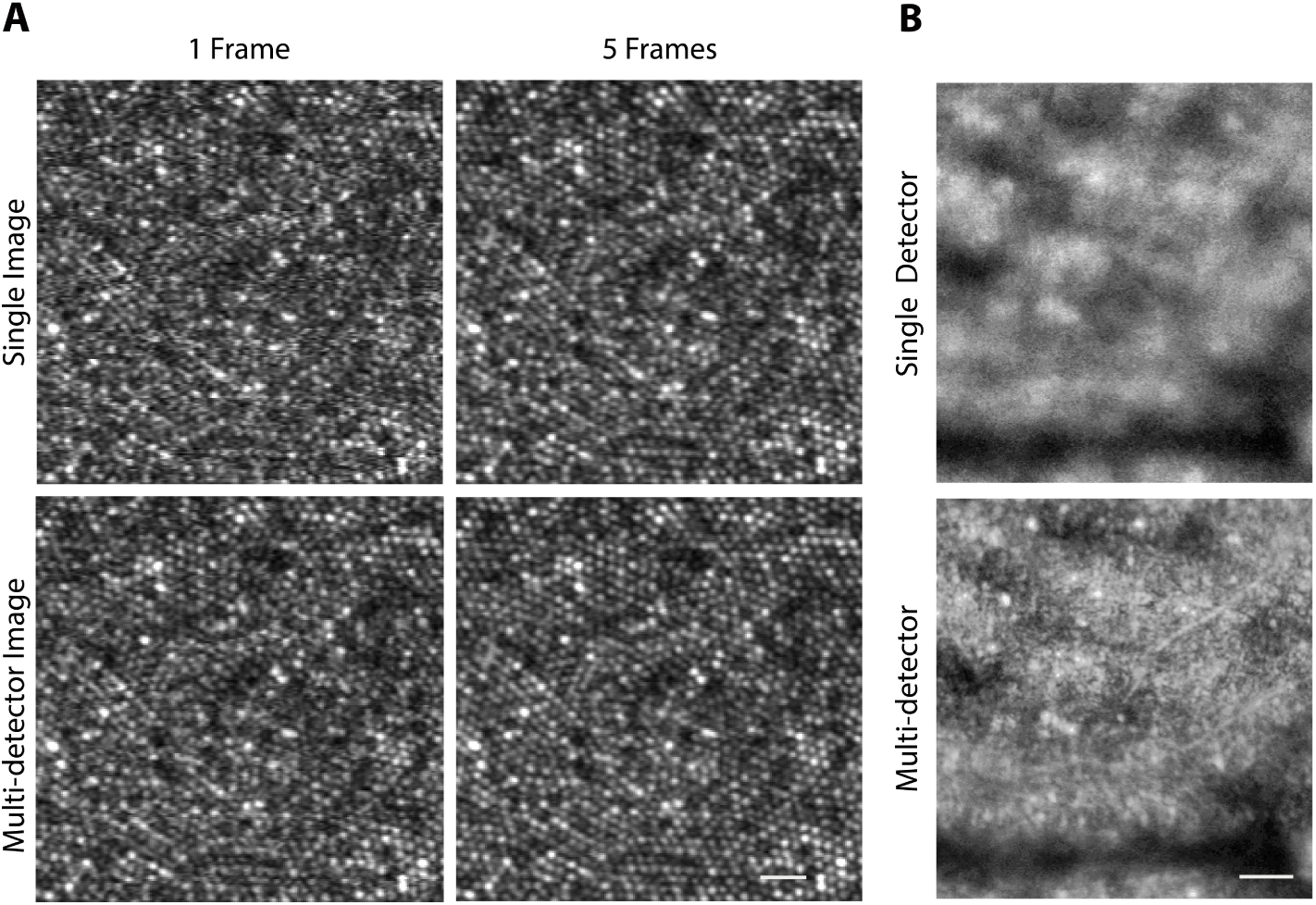
A. SNR analysis for signal to noise for a single channel on top row and the multi-detector on bottom row. B. The top image registers the eye movement stabilization from one imaging channel, the bottom image registers eye motion from the multi-detector channels exploiting the enhanced SNR. A. Subject 20112 Scale bar: 10*µm* B. Subject 20076 Scale bar: 20*µm*

The importance of SNR improvement within the acquisition time of a single frame becomes more evident in panel B of figure 5. These images were taken from a location in the retina 71.4*µm* superficial to the photoreceptor layer, where the retinal backscattered signal was weak. The SNR of the central channel alone was insufficient for strip-based frame registration, as can be seen in the upper image, which is blurred from uncorrected intra-frame image distortions. In the lower image, a higher SNR video using pixel reassignment was produced prior to applying the intra-frame eye movement correction algorithm. The resulting image has much higher frequency content and contrast due to the improved image registration accuracy.

The SNR was determined using the radial power spectrum information. The signal was quantified as the power spectrum value at the peak of the cone photoreceptor spatial frequency (∼125 cycles per degree) and the noise was quantified as the average signal above 140 cycles per degree, which is just beyond the peak signal expected from the photoreceptor mosaic. The plot on figure 6 shows that it takes about 3 times longer to reach the same level of SNR as the multi-detector scheme when imaging with a single detector.

**Fig. 6.**
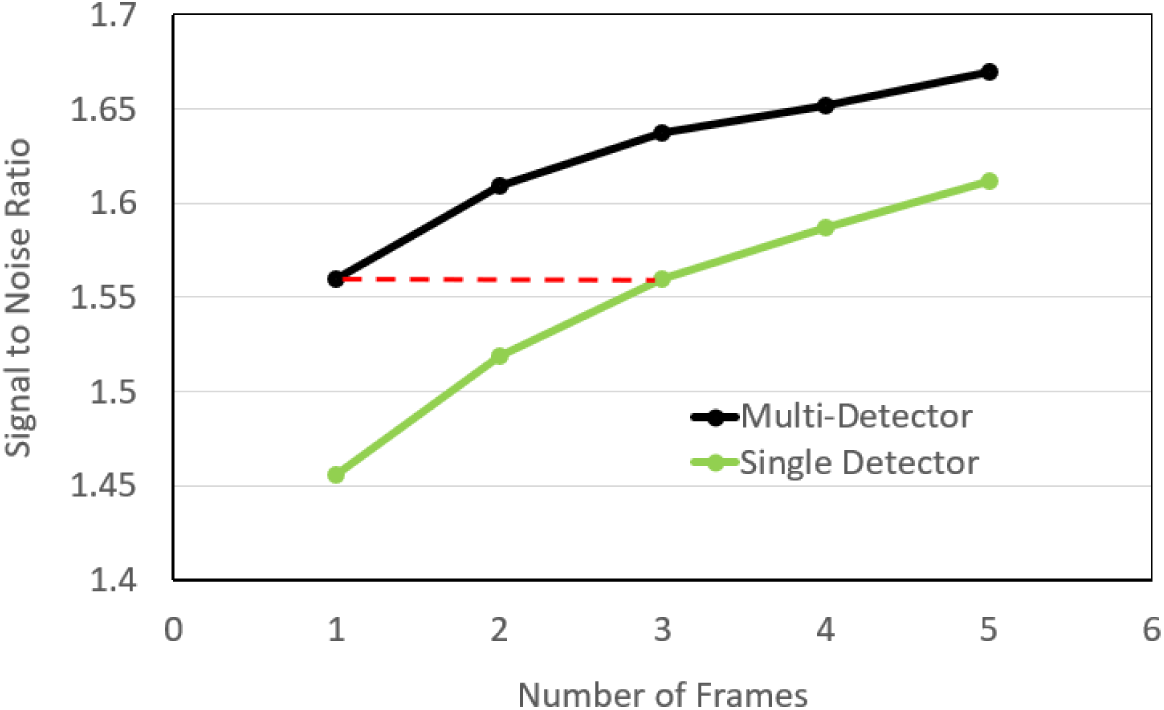
The SNR analysis shows that one multi-detector frame is equivalent in SNR to a three frame-average from the single detector as annotated in red.

### 3.2. Offset Aperture Imaging

In this configuration, the multi-detector telescope was set up to magnify the spot size by 0.15 to provide a collection PSF diameter of 13.8*µm*, which made the fiber core size become 14.5*ADD* for offset imaging. The optical power output of the imaging system prior to the eye was measured to be 108.6*µW* for 940*nm* and 73.6*µW* for 680*nm*. For figure 8 the 840*nm* imaging channel was used for eye motion registration with an optical power output of 52.3*µW*. First, three 10-second videos (900 frames) were collected for each individual detector and corrected for intra-frame eye motion.

Offset aperture imaging was able to reveal individual vascular layers and provide depth sectioning throughout the retina. Shown in figure 7 are the individual fiber images in which the center image is the confocal image and the outer images are from light collected from 13 to 19 ADD away from the center fiber via the outer fibers. To enhance the visualization of the refracted light by retinal structures, the differences of the opposing fibers were calculated and normalized by their sum. Here we can appreciate the different spatial orientations captured from each individual fiber. For example: fiber 2 shows the striations of the nerve fiber layer whereas fiber 6 shows horizontal processes emerging from the purported cell in the center of the image. In effect, each of the images from each fiber is tuned to retinal features of different orientations.

**Fig. 7.**
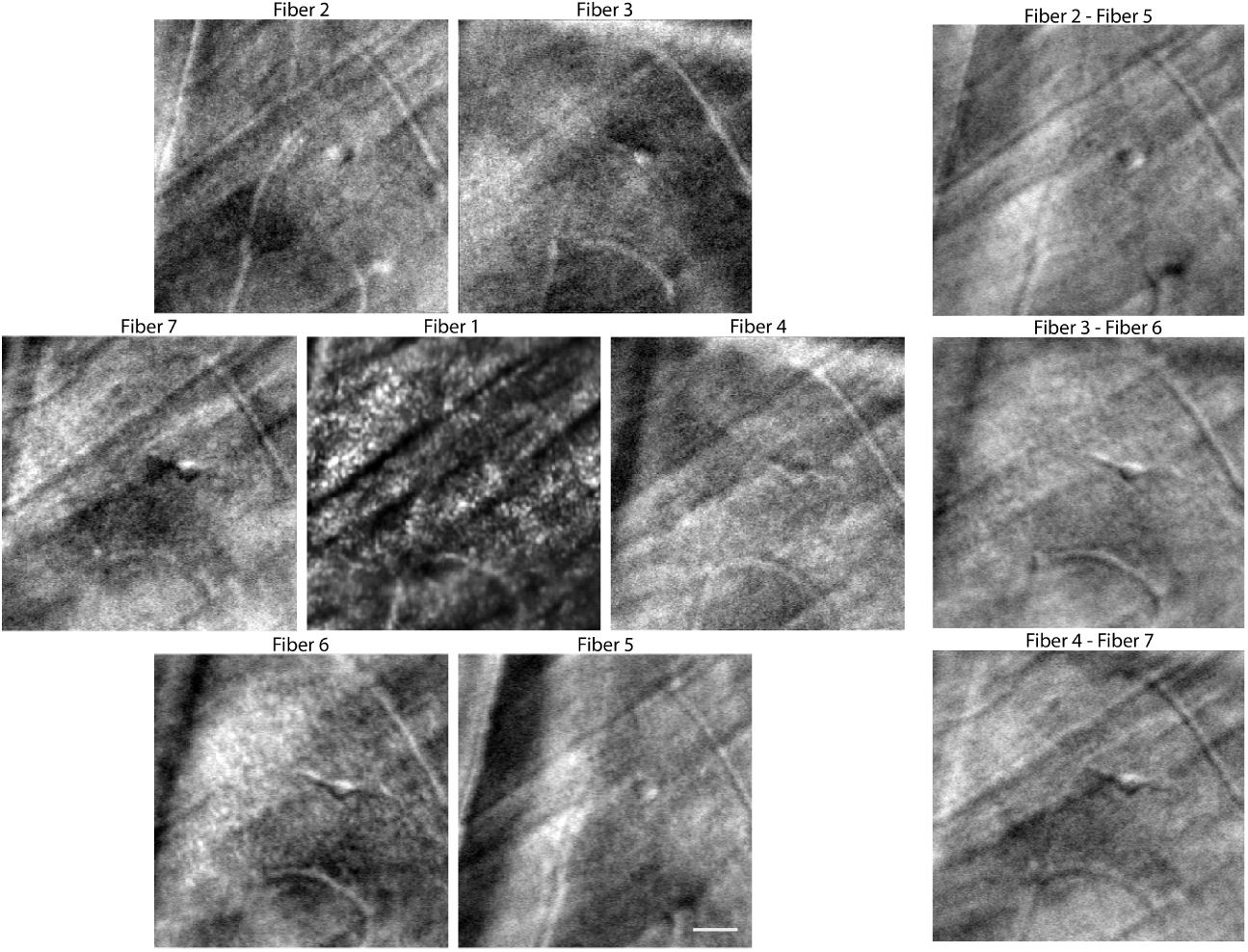
Left: The center image is the confocal image and the outer images are from light collected from areas between 13-19 ADD away from the confocal aperture. Right: Difference images from opposing fibers (normalized by their sum). Images were from Subject 20076. Scale bar: 30*µm*

In the process of testing the multi-detector scheme for offset aperture imaging, we frequently observed cells on the inner surface of the retina that had not been reported previously. We believe these are glial cells - either astrocytes or microglia. Figure 6 presents a closer examination of two of these cells in subject 20076. To confirm that these structures were not cross-sections of blood vessels running through the tissue, two analyses were performed. First, we did a perfusion analysis. The perfusion images, shown in the right column of figure 8, were generated by computing motion-contrast within stabilized videos [32]. In perfusion images, stable features appear dark while moving features (in this case, blood flow) appear white. The features indicated by green arrows are not apparent in the perfusion images, indicating that these structures are not likely to be blood vessels. Second, we did a through-focus analysis to rule out that the observed features were optical sections of longer, vessel-like structures. Two depth sections, separated by only 15 microns suggest that the features are isolated to a single layer in the retina. Although we expect microglia and astrocytes to be much more common in the human retina [33], it is possible that these are the only ones observed because of their superficial location. Similar cells residing within the retinal tissue might have a more similar refractive index to the surrounding tissue giving rise to a much smaller offset aperture signal.

**Fig. 8.**
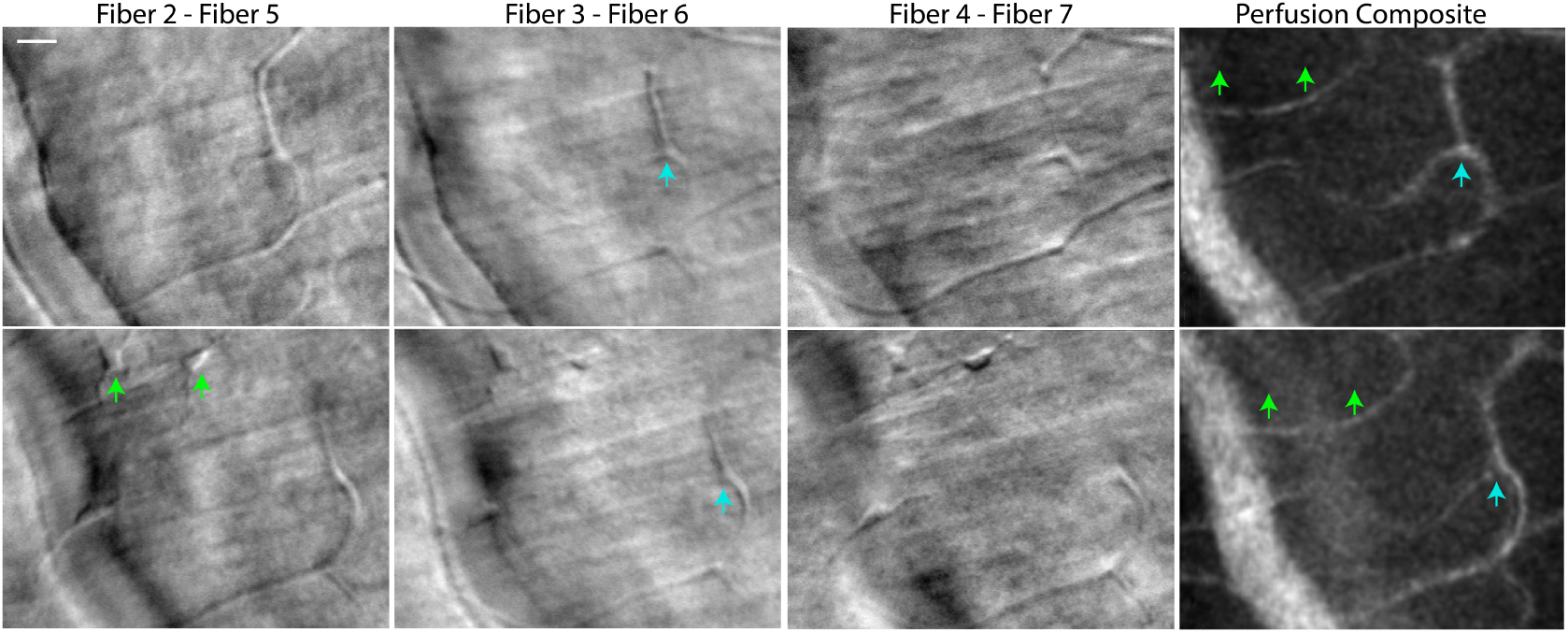
All images are taken at a depth location near the surface of the nerve fiber layer, the top row is at 15*µm* anterior (toward the vitreous) of the bottom row. The first three columns display difference images between opposing fibers. The last column shows perfusion maps of these areas to confirm that the structures indicated by the green arrows are not likely to be blood vessels. The blue arrow indicates a blood vessel for comparison. Images were acquired from subject 20076. Scale bar: 20*µm*

## 4. Discussion

We have demonstrated a versatile multi-detector system that uses a fixed fiber bundle detector array and enables either pixel reassignment or multi-offset aperture imaging.

To our knowledge, this is the first ever report of pixel reassignment in an AOSLO system. We believe that this detection modality, which offers optimal confocal resolution without compromising SNR, can be very useful in some imaging situations. Here is a short list of potential benefits for pixel-reassignment imaging.

1. Minimized phototoxicity: The maximum permissible exposures for visible wavelengths are very low, primarily to prevent photochemical damage to the retina [34]. However, the use of these wavelengths for imaging are useful for oximetry [35], fluorescence imaging (eg. fundus autofluorescence, fluorescein angiography) and other applications. Minimizing exposures without compromise to SNR or resolution will reduce the chances for incidental photo-toxic light exposures.

2. Better characterization of photopigments. Reduced powers at visible wavelengths will slow the bleaching rate of photoreceptors, improving the ability to measure photopigment absorption properties of individual cone photoreceptors, thereby making measurements of photopigment kinetics and AOSLO cone classing more efficient [36].

3. Reduced visibility of imaging beams. The scanning beam in current AOSLO systems are always visible. The visibility of the scanning raster can confound certain applications of functional testing in AOSLO systems, including microperimetry [37] and visual psychophysics [38, 39]. The ability to reduce light exposure and therefore visibility will be an important step towards invisible imaging.

4. Imaging weakly-reflective layers: Increased SNR will improve imaging and image registration from less reflective layers of the retina, eg inner and outer nuclear layers. Low backreflected signal makes these retinal structures difficult to analyze. Pixel reassignment both increases the signal throughput and enhances motion registration enabling a structural analysis of these inner retinal layers.

Our particular implementation of the multi-detector AOSLO scheme does come at a cost, which is lost light due to the void space between closely-packed fibers. Gaps between fibers (17% loss) and the cladding of the individual fibers (additional 17% loss) result in a total light loss of 34% compared to a circular aperture with the dimensions of the fiber bundle. The acceptance aperture of the multi-mode fiber is relatively large at 0.39 incurring only minimal losses. Nevertheless, the increase in signal with this scheme is still superior to that of a single detector that achieves the same level of resolution.

An attractive feature of the multi-detector scheme presented here is that it can be quickly reconfigured to different imaging modes by rearranging the same optical components to pre-determined locations. This offers versatility to the system, especially for a clinical system in which one might wish to acquire the highest possible resolution images at a specific location with high SNR and then utilize offset aperture imaging to probe the structural health of the transparent layers of the retina in the same patient. In this manuscript, we switched between two modes, which limited the range of offset aperture positions that could be explored. Images taken with the available offsets (13 - 19 ADD) images revealed intricate vascular information, inner retinal neurons and, for the first time in AOSLO, a few superficial cells speculated to be astrocytes or microglia. In future implementations, we intend to (i) add a zoom lens to replace lenses 1, 2 and 3 which will enable continuous control of detector offset positions, (ii) explore the use of different optical masks to fine-tune the size and shape of the offset detectors, and (iii) incorporate either an analog difference signal or use balanced detectors to minimize the noise floor and reduce the number of acquisition channels.

## 5. Conclusion

The multi-detector scheme is a versatile detection scheme enabling two different imaging configurations. The pixel reassignment configuration allows for more efficient light collection while preserving high spatial resolution, resulting in improved registration of natural eye movements in post-processing. The multi-offset imaging configuration reveals hidden phase structures such as blood vessels and individual cells.

## Acknowledgments

Supported by an Alcon Research Investigator Award (AR), the following grants from the National Institutes of Health (National Eye Institute): training grant: T32 EY007043 (SM), Bioengineering Partnership Grant: R01EY023591 (AR, VJ), Audacious Goals Initiative Grant: U01EY025501 (AR, FL, SM), Core grant: P30-EY003176 (AR, FL, PM, SM, VJ), Minnie Flaura Turner Memorial Fund for Impaired Vision Research (SM), Soroptimist International Founders Region Fellowship (SM)

## References and links

1. A. Roorda, F. Romero-Borja, W. J. Donnelly III, H. Queener, T. J. Hebert, and M. C. Campbell, “Adaptive optics scanning laser ophthalmoscopy,” Optics Express 10, 405 (2002).

2. T. Wilson and A. R. Carlini, “Size of the detector in confocal imaging systems,” Optics Letters 12, 227 (1987).

3. N. Sredar, O. E. Fagbemi, and A. Dubra, “Adaptive Optics Reflectance Confocal Scanning Light Ophthalmoscopy with Sub-Airy Disk Detectors,” Investigative Ophthalmology & Visual Science 58, 295 (2017).

4. A. Dubra and Y. Sulai, “Reflective afocal broadband adaptive optics scanning ophthalmoscope,” Biomedical Optics Express 2, 1757 (2011).

5. C. J. R. Sheppard, “Super-resolution In Confocal Imaging,” Optik 80, 53–54 (1988).

6. M. Defrise and C. De Mol, “Super-resolution in confocal scanning microscopy: gerenralized inversion formulae,” Inverse Problems 195 (1991).

7. M. Castello, C. J. R. Sheppard, A. Diaspro, and G. Vicidomini, “Image scanning microscopy with a quadrant detector,” Optics Letters 40, 5355 (2015).

8. J. Huff, “The Airyscan detector from ZEISS: confocal imaging with improved signal-to-noise ratio and super-resolution,” Nature Methods 12 (2015).

9. C. Roider, M. Ritsch-Marte, and A. Jesacher, “High-resolution confocal Raman microscopy using pixel reassignment,” Optics Letters 41, 3825 (2016).

10. D. Zhu, Y. Fang, Y. Chen, and A. Hussain, “Comparison of multi-mode parallel detection microscopy methods,” Optics Communications 387, 275–280 (2017).

11. C. B. Müller and J. Enderlein, “Image scanning microscopy,” Physical Review Letters 104, 1–4 (2010).

12. S. Roth, C. J. R. Sheppard, K. Wicker, and R. Heintzmann, “Optical photon reassignment microscopy (OPRA),” Optical Nanoscopy 2, 5 (2013).

13. G. M. R. De Luca, R. M. P. Breedijk, R. A. J. Brandt, C. H. C. Zeelenberg, B. E. de Jong, W. Timmermans, L. N. Azar, R. A. Hoebe, S. Stallinga, and E. M. M. Manders, “Re-scan confocal microscopy: scanning twice for better resolution,” Biomedical Optics Express 4, 2644 (2013).

14. T. Azuma and T. Kei, “Super-resolution spinning-disk confocal microscopy using optical photon reassignment,” Optics Express 23, 15003 (2015).

15. A. G. York, S. H. Parekh, D. D. Nogare, R. S. Fischer, K. Temprine, M. Mione, A. B. Chitnis, C. A. Combs, and H. Shroff, “Resolution doubling in live, multicellular organisms via multifocal structured illumination microscopy,” Nature Methods 9, 749–754 (2012).

16. A. G. York, P. Chandris, D. D. Nogare, J. Head, P. Wawrzusin, R. S. Fischer, A. Chitnis, and H. Shroff, “Instant super-resolution imaging in live cells and embryos via analog image processing,” Nature Methods 10, 1122–1130 (2013).

17. O. Schulz, C. Pieper, M. Clever, J. Pfaff, A. Ruhlandt, R. H. Kehlenbach, F. S. Wouters, J. Grosshans, G. Bunt, and J. Enderlein, “Resolution doubling in fluorescence microscopy with confocal spinning-disk image scanning microscopy,” Proceedings of the National Academy of Sciences 110, 21000–21005 (2013).

18. F. LaRocca, T. B. DuBose, S. Farsiu, and J. A. Izatt, “Optical photon reassignment super-resolved scanning laser ophthalmoscopy (Conference Presentation),” Investigative Ophthalmology & Visual Science 58, 100451C (2017).

19. A. Elsner, S. A. Burns, J. J. Weiter, and F. C. Delori, “Infrared imaging of sub-retinal structures in the human ocular fundus,” Vision Research 36, 191–205 (1996).

20. D. Scoles, Y. N. Sulai, and A. Dubra, “In vivo dark-field imaging of the retinal pigment epithelium cell mosaic,” Biomedical Optics Express 4, 1710 (2013).

21. D. Scoles, Y. N. Sulai, C. S. Langlo, G. A. Fishman, C. A. Curcio, J. Carroll, and A. Dubra, “In Vivo Imaging of Human Cone Photoreceptor Inner Segments,” Investigative Opthalmology & Visual Science 55, 4244 (2014).

22. T. Y. P. Chui, D. a. VanNasdale, and S. a. Burns, “The use of forward scatter to improve retinal vascular imaging with an adaptive optics scanning laser ophthalmoscope,” Biomedical Optics Express 3, 2537 (2012).

23. A. Guevara-Torres, A. Joseph, and J. B. Schallek, “Label free measurement of retinal blood cell flux, velocity, hematocrit and capillary width in the living mouse eye,” Biomedical Optics Express 7, 4228 (2016).

24. A. Guevara-Torres, D. R. Williams, and J. B. Schallek, “Imaging translucent cell bodies in the living mouse retina without contrast agents,” Biomedical Optics Express 6, 2106 (2015).

25. E. A. Rossi, C. E. Granger, R. Sharma, Q. Yang, K. Saito, C. Schwarz, S. Walters, K. Nozato, J. Zhang, T. Kawakami, W. Fischer, L. R. Latchney, J. J. Hunter, M. M. Chung, and D. R. Williams, “Imaging individual neurons in the retinal ganglion cell layer of the living eye,” Proceedings of the National Academy of Sciences 114, 586–591 (2017).

26. K. Sapoznik, T. Luo, R. L. Warner, A. De Castro, L. Sawides, and S. A. Burns, “Enhanced imaging of retinal vessels using a configurable aperture AOSLO,” Investigative Ophthalmology & Visual Science 58, 3434 (2017).

27. K. Grieve, P. Tiruveedhula, Y. Zhang, and A. Roorda, “Multi-wavelength imaging with the adaptive optics scanning laser Ophthalmoscope,” Optics Express 14, 12230 (2006).

28. C. T. Rueden, J. Schindelin, M. C. Hiner, B. E. DeZonia, A. E. Walter, E. T. Arena, and K. W. Eliceiri, “ImageJ2: ImageJ for the next generation of scientific image data,” BMC Bioinformatics 18, 1–26 (2017).

29. S. B. Stevenson, A. Roorda, and G. Kumar, “Eye tracking with the adaptive optics scanning laser ophthalmoscope,” Proceedings of the 2010 Symposium on Eye-Tracking Research & Applications - ETRA ’10 p. 195 (2010).

30. I. J. Cox, C. J. R. Sheppard, and T. Wilson, “Improvement in resolution by nearly confocal microscopy,” Applied Optics 21, 778 (1982).

31. M. Guizar-Sicairos, S. T. Thurman, and J. R. Fienup, “Efficient subpixel image registration algorithms.” Optics letters 33, 156–158 (2008).

32. J. Tam, J. A. Martin, and A. Roorda, “Noninvasive visualization and analysis of parafoveal capillaries in humans,” Investigative Ophthalmology and Visual Science 51, 1691–1698 (2010).

33. T. Chan-Ling, “Glial, neuronal and vascular interactions in the mammalian retina,” Progress in Retinal and Eye Research 13, 357–389 (1994).

34. J. I. W. Morgan, J. J. Hunter, B. Masella, R. Wolfe, D. C. Gray, W. H. Merigan, F. C. Delori, and D. R. Williams, “Light-induced retinal changes observed with high-resolution autofluorescence imaging of the retinal pigment epithelium,” Investigative Ophthalmology and Visual Science 49, 3715–3729 (2008).

35. H. Li, J. Lu, G. Shi, and Y. Zhang, “Measurement of oxygen saturation in small retinal vessels with adaptive optics confocal scanning laser ophthalmoscope,” Journal of Biomedical Optics 16, 110504 (2011).

36. R. Sabesan, H. Hofer, and A. Roorda, “Characterizing the human cone photoreceptor mosaic via dynamic photopigment densitometry,” PLoS ONE 10, e0144891 (2015).

37. W. S. Tuten, P. Tiruveedhula, and A. Roorda, “Adaptive optics scanning laser ophthalmoscope-based microperimetry,” Optometry and Vision Science 89, 563–574 (2012).

38. R. Sabesan, B. P. Schmidt, W. S. Tuten, and A. Roorda, “The elementary representation of spatial and color vision in the human retina,” Science Advances 2, e1600797 (2016).

39. N. Domdei, L. Domdei, J. L. Reiniger, M. Linden, F. G. Holz, A. Roorda, and W. M. Harmening, “Ultra-high contrast retinal display system for single photoreceptor psychophysics,” Biomedical Optics Express 9, 157 (2018).

